# Resolving heterogeneous high-mass macromolecular machineries by Orbitrap-based single particle charge detection mass spectrometry

**DOI:** 10.1101/717413

**Authors:** Tobias P. Wörner, Joost Snijder, Antonette Bennett, Mavis Agbandje-McKenna, Alexander A. Makarov, Albert J.R. Heck

## Abstract

Here we show that single particle charge-detection mass spectrometry (CD-MS) can be performed on a ubiquitous Orbitrap mass analyser and applied to the analysis of high-mass (megadalton) heterogeneous biomolecular assemblies. We demonstrate that single particle high-mass ions can survive in the Orbitrap for seconds, whereby their measured signal amplitudes scale linearly with charge state over the entire *m/z* range. Orbitrap based single particle CD-MS can be used to resolve mixed ion populations, accurately predict charge states, and consequently also the mass of the ions. We successfully applied CD-MS to challenging natural and biotherapeutic protein assemblies, such as IgM oligomers, designed protein nano-cages, ribosome particles and intact, empty- and genome-loaded Adeno-associated virus particles. Single particle CD-MS combined with native MS on existing Orbitrap platforms will greatly expand its application, especially in the mass analysis of megadalton heterogeneous biomolecular assemblies.

## Main text

Over the past two decades, native mass spectrometry (MS) has developed into a powerful tool for the characterization of biological macromolecular complexes ^1^. These complexes are analysed from solvents that preserve the noncovalent interactions, enabling mass analysis of intact macromolecular assemblies into the megadalton (MDa) range ^2^. The exact mass of the intact macromolecular complex is then used to infer its composition, the subunit stoichiometry, and even the number and chemical nature of post-translational modifications and small molecule ligands bound to the complex ^3–6^. Various mass analysers, including Quadrupole Time-of-Flight, Fourier-Transform Ion Cyclotron Resonance, and most recently Orbitrap™-based mass analysers, have all been adapted for native MS experiments ^7–12^. With the successful modifications of the Orbitrap for use in native MS, an unprecedented mass resolving power in the high mass range could be achieved, opening the avenue for high resolution analysis on ever more heterogeneous protein assemblies like therapeutic and plasma glycoproteins, designed protein cages, membrane protein assemblies, intact viruses, and ribosomes ^12,13^.

Notably, masses are not measured directly in most MS approaches, but need to be inferred from the mass-to-charge (*m/z*) ratios of the detected ions. As first shown in the pioneering work of Mann and Fenn ^14^, the charge states from a population of multiply charged ions generated by electrospray ionization (ESI) can be determined from the *m/z* values by matching consecutive peaks in the charge state distribution to calculate accurate masses. A general limitation in native MS studies then stems from the fact that the charge state, and thus also the mass, can only be accurately measured when multiple charge states of the same molecular species can be resolved and assigned in the *m/z* spectrum. This hampers the analysis of larger heterogeneous protein assemblies, such as amyloid fibrils, genome packed viruses, and membrane protein complexes decorated with multiple lipid molecules.

Even small variabilities in monomeric building blocks, commonly originating from protein N-or C-terminal truncations and other post-translational modifications, can result in wide distributions of masses in larger assemblies with a high number of subunits. In combination with the often-poor desolvation of large assemblies, these broadened mass distributions result in overlapping signals between consecutive charge states, leading to inaccurate mass assignments. For instance, when charge states cannot be resolved, masses are sometimes still estimated from the centroid *m/z* of a broadly distributed signal, assuming a normal charging behaviour for globular proteinaceous particles ^15,16^. But when particles don’t show normal charging behaviour, for instance when considerable amounts of RNA or DNA are present in the complex ^12^, this approach may yield false mass assignments.

An attractive alternative may come from measuring one particle (or ion) at a time, thereby avoiding the problematic convolution of signals that stem from insufficient resolving power ^17–27^. When such single-particle detection approaches can be combined with an independent measure of the charge of ions, or when masses can be estimated using entirely different physical principles that circumvent the need to work with multiply charged ions, this opens up the door to bona fide single-particle mass spectrometry measurements.

Several techniques for single particle mass spectrometry, capable of determining masses into the MDa range have emerged in recent years, most notably Nano Electro-Mechanical System mass spectrometry (NEMS-MS) and Charge Detection mass spectrometry (CD-MS) ^21,22^. In NEMS-MS, single particles are deposited on a NEMS resonator, whose vibrational frequency is affected by the absorption of a single particle. The mass of the impacting particle can be calculated directly from the frequency shift that the resonator experiences ^19,23–25^. In CD-MS, the mass is calculated by measuring the *m/z* ratio and charge separately, typically by oscillating the particle in a cone trap through a conductive detection tube ^17,26,27^. Knowing the ion energy, the charge can be determined from the pulse amplitude of the image current induced by the particle passing through and the *m/z* can be derived from the frequency of the oscillation. Both NEMS-MS and CD-MS approaches have demonstrated their capability in the analysis of large biomolecular complexes, especially of viruses in the 1-100 MDa range ^27–32^.

Ion transmission has usually been a limiting factor in native MS and great efforts in instrument design have been made to transmit sufficient ions up to the MDa range, which often still requires hours of signal accumulation ^9–12^. However, the Orbitrap mass analyser is sensitive enough for the detection of single (multiply charged) ions, opening up the possibility for Orbitrap-based single particle mass analysis ^11,33,34^. It should be noted that single ion detection was demonstrated much earlier on FT-ICR instruments ^35^. Following up the early work by Makarov *et al.* on myoglobin sprayed under denaturing conditions ^33^, we demonstrated single ion detection in native MS for the 800 kDa GroEL complex with approximately 70 charges ^11^. More recently, Kafader *et al.* reported^34^ that by centroiding, binning, and stringent filtering of the signals of individual ions, it is possible to enhance the Orbitrap instrument’s effective resolution by an order of magnitude compared to measurements on ion ensembles, which allowed the authors of (34) to resolve the isotopic distribution of intact antibodies sprayed under denaturing conditions.

Here, we demonstrate that the intensity of a single ion detected in an Orbitrap can be directly used to infer its charge state, thus enabling single particle mass spectrometry. The recorded signal in the Orbitrap analyser corresponds to the image current induced by the oscillating ions and we show here that its amplitude scales linearly with the ion’s charge. We examined single ions over a very wide *m/z* and charge range for several proteins and protein assemblies, ranging from 150 kDa to 9.4 MDa in mass, empirically establishing a highly correlated linear dependency between intensity and charge. Importantly for the work here, we noticed that these megadalton-sized single ions are extremely stable, allowing us to record image current signals of more than a second. Subsequently, we used Orbitrap-based charge detection mass spectrometry to distinguish and determine the mass of several co-occurring IgG and IgM multimers, demonstrating that the intensity of single ions can be used as an additional dimension to resolve otherwise overlapping charge state distributions. Furthermore, we demonstrate that by using Orbitrap-based CD-MS charge states can be estimated from single ion intensities, enabling mass measurements of single particles. These masses determined by CD-MS are in excellent agreement with native MS measurements on ion ensembles based on conventional charge state assignments.

The development of Orbitrap-based CD-MS opens up analysis of a whole new range of macromolecular assemblies that would be inaccessible to conventional native MS due to their mass heterogeneity. As an example of such applications, we analysed a mixture of empty and genome filled Adeno-associated virus (AAV) particles, currently one of the most favoured vectors responsible for the renaissance of gene-therapy applications ^36,37^. Orbitrap-based CD-MS allowed us to not only distinguish empty from genome packed particles, but also to obtain accurate masses of both the empty capsid and the packed virus, providing a means to quality control genome packaging.

## Results

### Detection of single ions on the Orbitrap

As shown previously, the Orbitrap mass analyser is sensitive enough to detect single multiply charged ions ^11,33,34^. During acquisition, ions get injected from the C-trap into the Orbitrap analyser and start oscillating with a radial, angular and axial frequency. Oscillation of the ions induces an image current on the outer electrodes (Fig. 1A). This image current is amplified and recorded over the duration of a set scan time, after which the frequency and amplitude of the oscillating signal is determined using a Fourier transformation of the transient signal to yield the final *m/z* spectrum. The number of charges on an ion determine the amplitude of the induced image current and thereby the intensity of the signal for the corresponding frequency in the final mass spectrum. Consequently, the signal intensities can be used as a proxy of ion abundances, which is a common assumption in quantitative MS measurements. By the same principle, the signal amplitude of a single ion should be a useful proxy to estimate the number of charges it carries, independent of the frequency (*i.e. m/z*) of the signal. If the scaling of the absolute signal intensities with the number of charges can be determined, this opens the door to single-particle CD-MS on an Orbitrap mass analyser, as both *m/z* and charge can be estimated for every individual ion.

**Figure 1:**
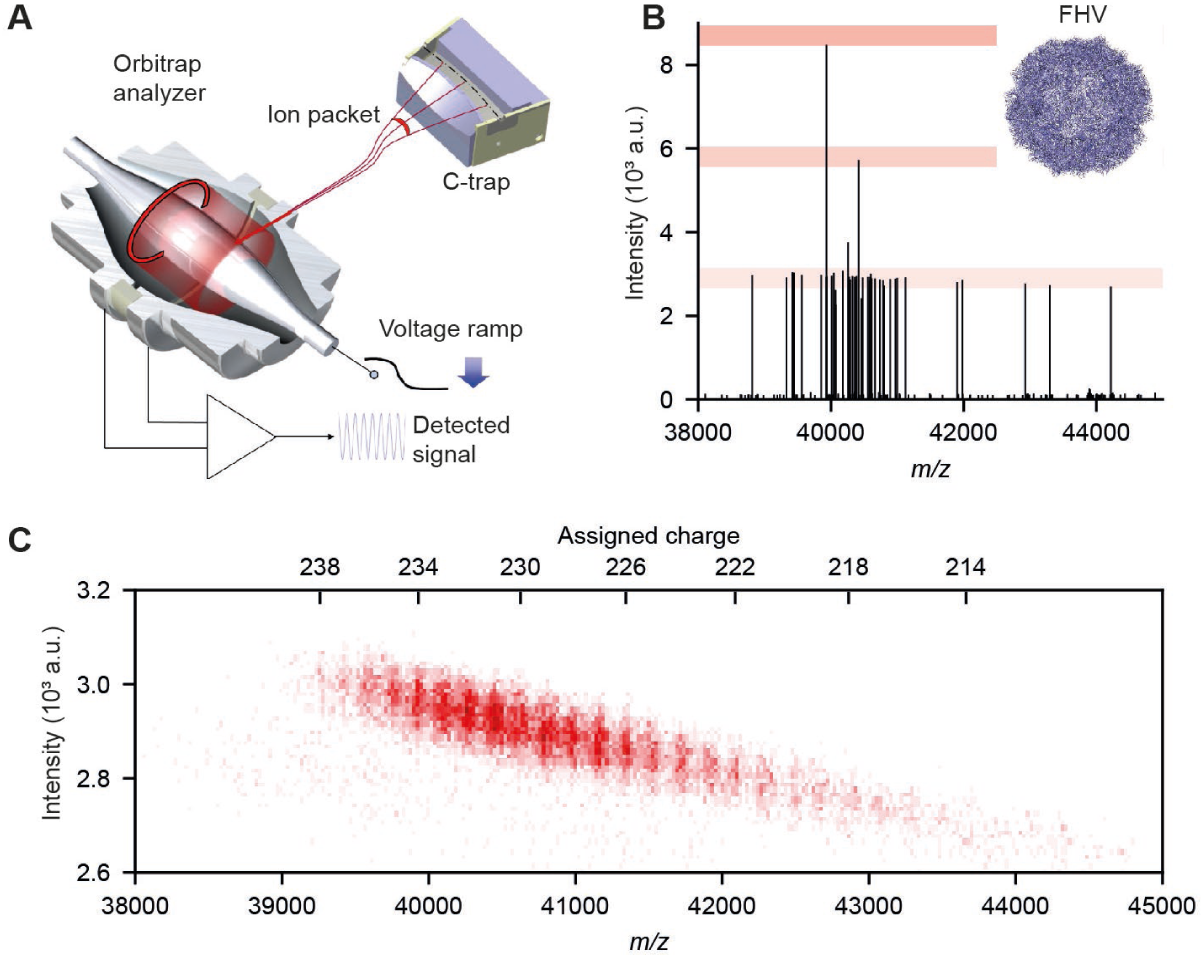
Single ion detection with the Orbitrap mass analyser. **A** Schematic of the injection of ions from the C-trap into the Orbitrap, where they oscillate along the central electrode, inducing an image current, which is recorded and converted into a final m/z spectrum by Fourier transformation. **B** A single scan showing several dozens of individual multiply charged ions of the Flock house virus (FHV) with discrete intensities around 3*10^3^, highlighted by the bottom red bar. Two instances are visible where two and three ions coincide at the same frequency (m/z), yielding two- and three-fold intensities of the single ions, highlighted by the middle and top red bars, respectively. **C** 2D histogram of filtered single ion signals collected over several minutes of acquisition time. The bottom x-axis indicates the m/z, whereas the top x-axis indicates the charge states. Note that the signal intensity of single FHV particles decreases with decreasing charge over the entire displayed region.

To achieve experimentally the storage and thus subsequent detection of single ions in an Orbitrap, we had to dilute analytes to the nanomolar range, shorten the time during which ions are sampled from the continuous beam produced by ESI to the order of milliseconds, or down-tune the ion-optics to further reduce ion transmission. When the sample of ions becomes sufficiently attenuated, a spectrum from a single scan will contain widely distributed sharp peaks and discrete signal intensities for ions with similar charge states. Additionally, if multiple ions of the same charge coincide at a given *m/z* within the same scan, their combined intensity will appear as an integer multiple of the corresponding single ion’s intensity. This is illustrated in Fig. 1B, where we show a single scan recorded for a sample of flock house virus (FHV) particles. Many ions are detected simultaneously, though most ions have a unique *m/*z in the region around 41,000 *m/z*, with approximately 220 charges. With such a relatively high number of charges, the single ion signals can be easily distinguished against noise (S/N ∼ 250). The small spread of single ion intensities for individual charges confirms that most ions survive until the end of the transient. The average lifetime of such ions in the Orbitrap therefore extends to at least several seconds. This somewhat surprising stability of high-mass ions accelerated by almost 2,000 V is in striking contrast to abundant fragmentation observed for smaller proteins at a better vacuum ^33^. Following (33), this nevertheless could be explained by reduction of centre-of-mass collision energy with molecules of residual gas (nitrogen or xenon) to below 2 eV, i.e. below fragmentation threshold of all covalent bonds.

Additionally, we observe a few incidences where two and tree ions coincide at the same *m/z*, yielding two- and three-fold intensities of single ions. Of note, the resolution on the single ion peaks closely follows the theoretical resolution limit of the instrument (Fig. S1). Whereas a multitude of ions is detected in every scan, almost all ions in the sample have a unique *m/z* and are thus detected and resolved as single ions. The multiplicity of ions in all scans substantially improves the duty cycle compared to measurements of a single ion at a time, as has been noted for other single particle MS approaches ^19^. With the ion sampling described here, thousands of ‘single ions’ can be detected within minutes. The ion intensities and *m/z* of thousands of FHV particles accumulated over a minutes-long measurement are displayed in Fig. 1C as a two-dimensional histogram. For this relatively homogeneous sample, charge states could be resolved and charges are indicated at the top x-axis. A careful assessment reveals a continuous decrease of single particle intensities with decreasing charges over the whole FHV charge state distribution, highlighting the dependency of charge and intensity within a single charge state distribution.

### Single ion amplitudes correlate linearly with charge state

To determine quantitatively how single ion intensities scale with the number of charges, we conducted single ion measurements on a larger set of protein complexes with well-defined masses, ranging from 150 kDa to 9.4 MDa (see Fig. 2A). Each sample was sprayed from aqueous ammonium acetate, with and without the addition of the charge reducing agent triethyl ammonium acetate (TEAA) ^38^. The use of TEAA allows us to additionally obtain single ion intensities for relatively low charges at high *m/z*, which improves sampling across the charge dimension and provides validation that the single ion intensity is not affected by the frequency (*m/z*) of the ions. The single ion events were binned by *m/z* after filtering detected single ion centroids for noise and decaying ion signals (see Fig. S4), yielding charge state resolved spectra that could be used for conventional charge state assignment. The assigned charge states were then compared to the single ion intensities at the corresponding *m/z* to build a calibration curve of single ion intensity vs charge state (see Fig. S5). This data is depicted in Fig. 2B, which shows 200 randomly sampled ions per charge state and a linear regression model describing the single ion intensity as a function of charge with an r^2^ of 0.997.

**Figure 2:**
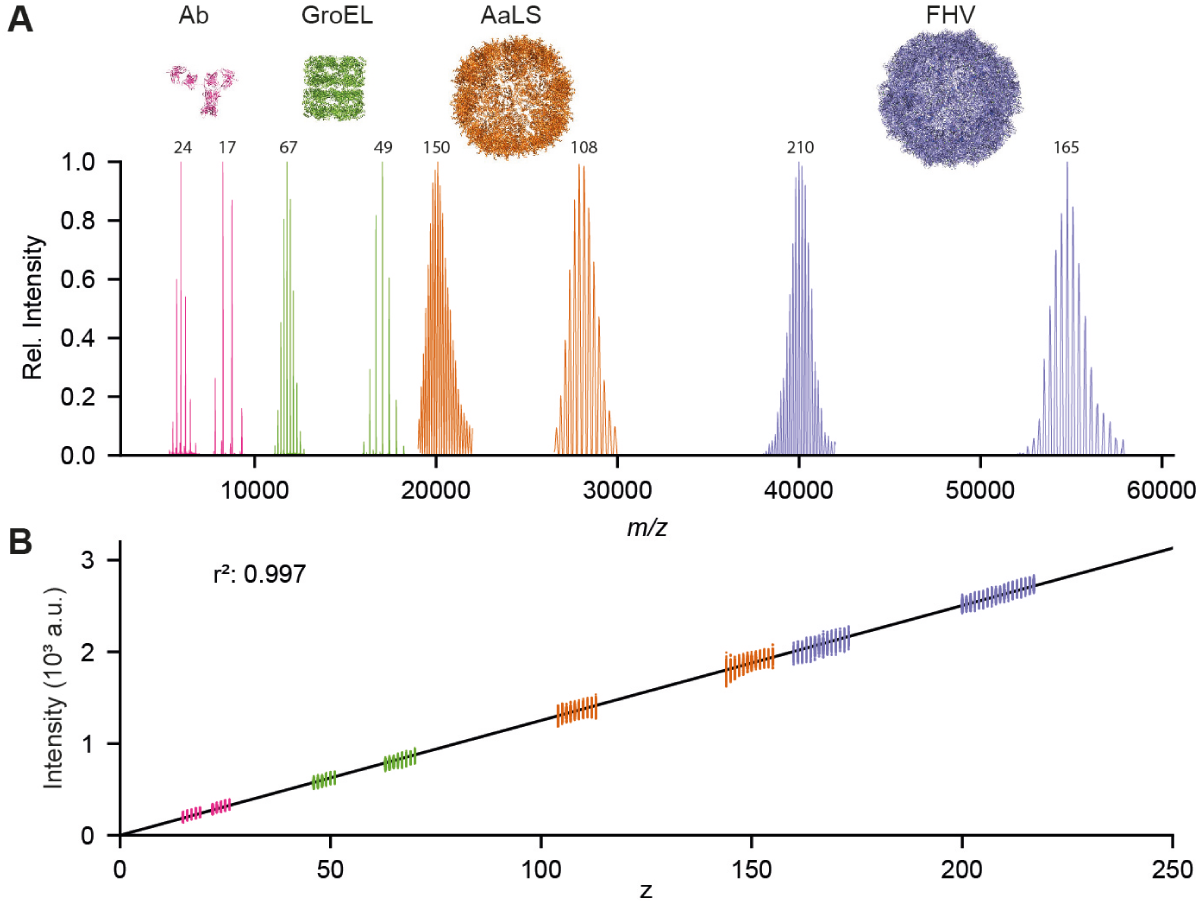
Single ion intensities scale linearly with charge state. **A** Composite native MS spectra of protein assemblies measured to evaluate the scaling of single ion intensity with charge state. Each protein assembly was analysed with and without the addition of the charge reducing agent TEAA in the electrospray solution, resulting for each species in two mass spectra at lower and higher m/z, respectively. The average charge state is indicated above each spectrum. **B** A linear regression model was fitted to 200 sampled single ion intensities per charge state: Intensity = 12.521 * charge, with an r^2^ of 0.997.

Using this linear relationship to predict the charge state of individual ions yields an RMSD of 3.5 charges. Based on the peak intensity of the combined ion populations sampled for the fit (as calculated in Fig. S5C) the RMSD is 1.2 charges, amounting to an error of less than 1% for the particles in the MDa range. This error is also reflected when the mass is predicted for each ion individually, with average masses for FHV and AaLS-neg not deviating more than 1.1% from the mass calculated by using the conventional charge state assignment (see Tab. S1). We performed a second calibration of intensity vs charge over the course of three months on the same instrument to test the stability of the single ion response on the detector. The resulting regression model of the second calibration is nearly identical with an r^2^ of 0.998 (see Fig. S2B). The maximum deviation between both calibrations does not exceed 1.6 charges in the range up to 250 charges, illustrating the excellent robustness of the established relationship between intensity and charge on a given Orbitrap mass analyser.

### Single particle analysis resolves mixed ion populations

Next, we tested how the Orbitrap mass analyser is capable of resolving more complex mixtures in the intensity domain. We used AaLS-neg, an engineered protein cage with a mass of 3 MDa, and the 50S ribosomal subunit of *E. coli* with a mass of 1.4 MDa. Both assemblies populate the same *m/z* region around 20,000 to 22,000 under native MS conditions. Since the 50S ribosomal particles contain around 67% RNA ^12^, it acquires much fewer charges during ESI compared to the protein assembly AaLS-neg, which we hypothesized should be reflected by the single ion intensities. An exemplary spectrum, and a 2D histogram of ion counts over a longer acquisition time are shown in Fig. 3A and Fig. 3B, respectively.

**Figure 3:**
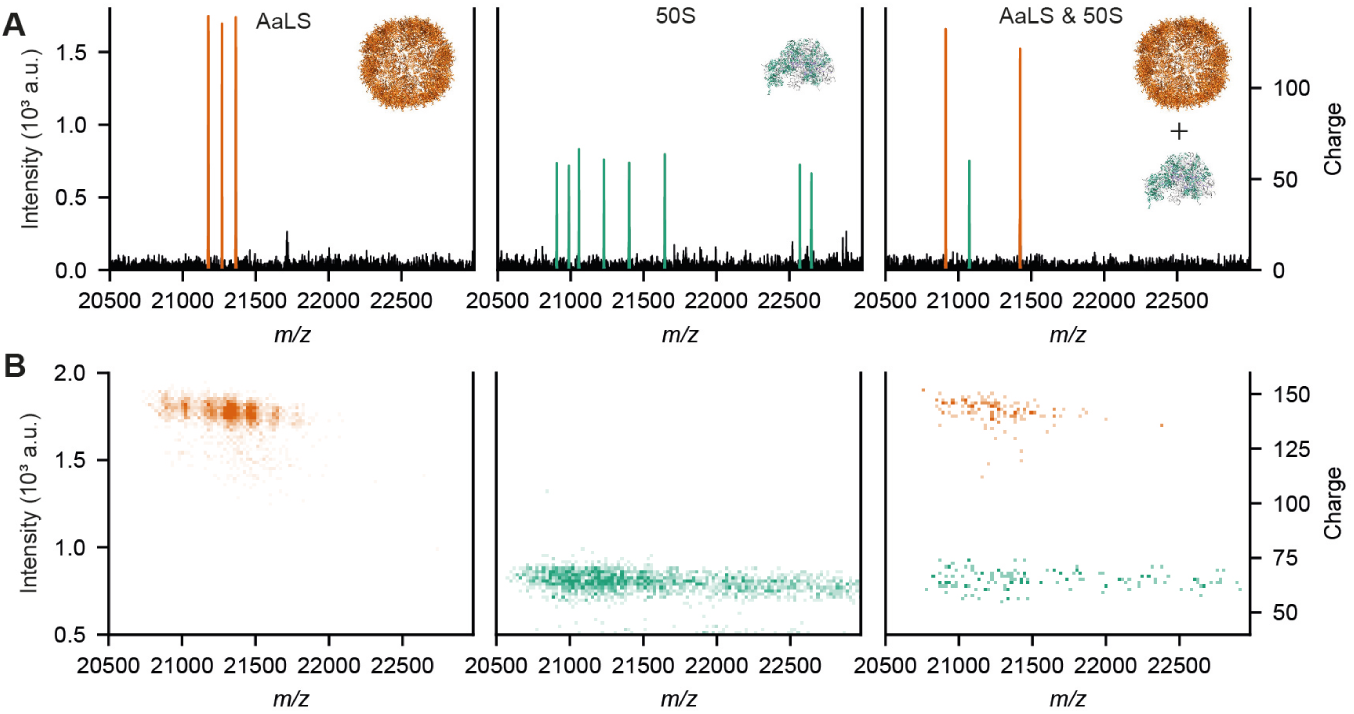
Resolving mixed ion populations with overlapping m/z based on single ion intensities. **A** Individual scans of single particles of AaLS-neg (left), 50S E. coli ribosomal particles (middle) (3.05 and 1.39 MDa, respectively) and a mixture of both these particles (right). Ion intensities display the expected intensity of ≈1.8*10^3^ for AaLS-neg and ≈0.8*10^3^ for the 50S particles. **B** Corresponding binned 2D histogram of larger datasets of single particles. Single ion signals can be easily assigned to AaLS-neg or 50S based on intensity, even within the same scan.

For AaLS-neg, we detect ions of the charge states 140-150 in the region between 21,000 *m/z* and 21,500 *m/z*. The 50S ribosomal subunit ions carrying 60-70 charges populate the same *m/z* region. Whereas the AaLS-neg and 50S ions overlap in *m/z*, the complexes can be clearly distinguished based on the single ion intensities, even when the two complexes are sprayed from a mixture and co-occur within the same scan. Hence, single ion intensities can be used to resolve mixed ion populations of the same *m/z*.

From this artificial mixture of proteins assemblies, we turned to naturally occurring complex mixtures, focusing first on the immunoglobulin variant IgG1-RGY. This is an IgG variant with three designed mutations that promote oligomerization in solution, with a preferred stoichiometry of six IgG molecules. This IgG oligomerization dramatically enhances its binding to the complement factor C1 ^39^. The resulting conventional native MS spectrum (see Fig. 4A) yields a wide range of charge states for each of the different oligomeric species (from monomer to hexamer), whereby in particular the charge state distributions of the trimeric and tetrameric species overlap in the *m/z* domain to a great extent. Although most charge states can be resolved on the QE-UHMR, charge states 42 for the trimer and 56 for the tetramer precisely coincide at 10,649 *m/z* (see Fig. 4B). However, the single ion intensities of the IgG oligomers scale with charge state (see Fig. 4C), and the trimeric and tetrameric signals at 10,649 *m/z* are completely resolved in this dimension.

**Figure 4:**
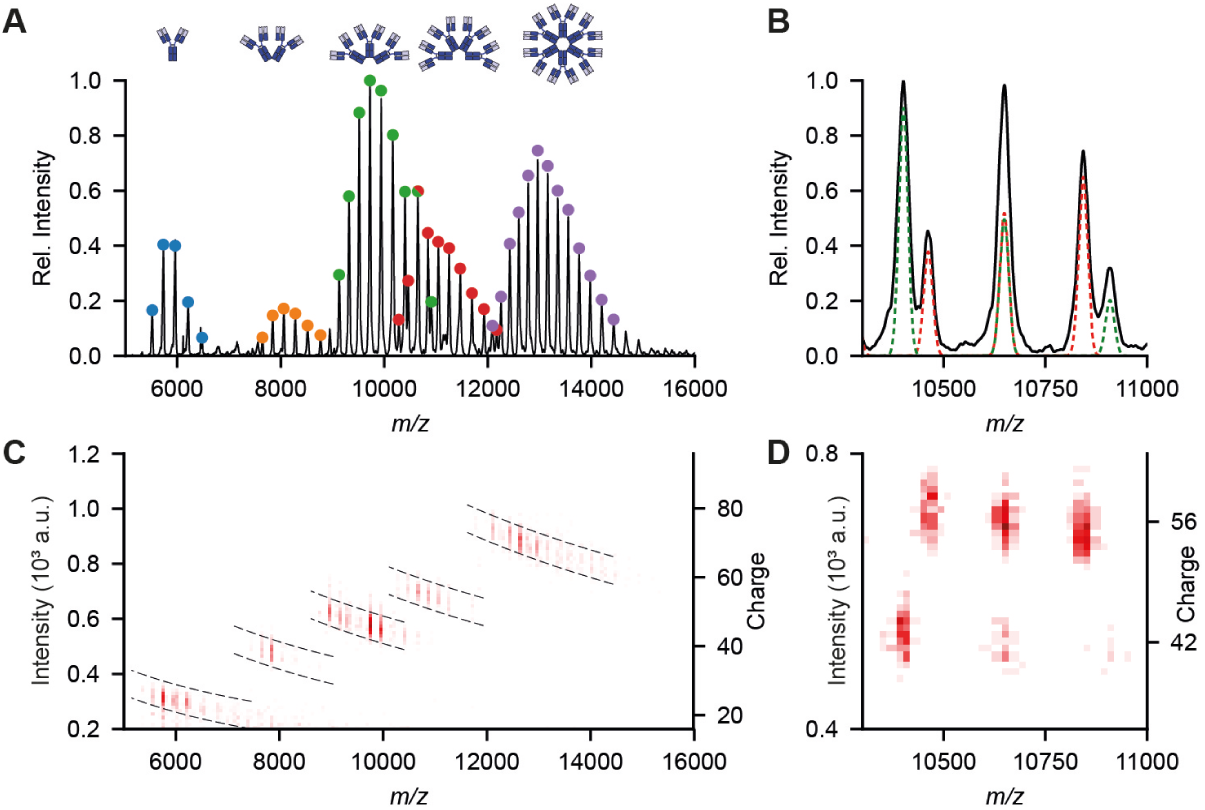
Resolving IgG oligomers based on single ion intensities. **A** Conventional native MS spectrum of IgG1-RGY. **B** Detailed view of the spectrum in A, showing the exact overlap between two charge states from the trimer (42+) and tetramer (56+) distributions at 10,649 m/z. **C** Single particle CD-MS spectrum of IgG1-RGY. **D.** Detailed view of the same region as in B, showing complete resolution of the coinciding trimer and tetramer charge states in the intensity dimension.

We next analysed IgM, which is thought to naturally exist in two co-occurring variants (pentamer and hexamer) ^40^. When we measured an IgM sample, recombinantly expressed without the J-chain, the Orbitrap based CD-MS spectra clearly revealed signals from three distinct co-occurring oligomeric species. The multiple occupied glycosylation sites on IgM introduce a high grade of heterogeneity, which cannot be resolved at this high *m/z* region causing broad peaks for each individual charge state. Additionally, the three charge state distributions overlap extensively, resulting in a poorly resolved complex spectrum and preventing accurate charge state assignment (see Fig. 5).

**Figure 5:**
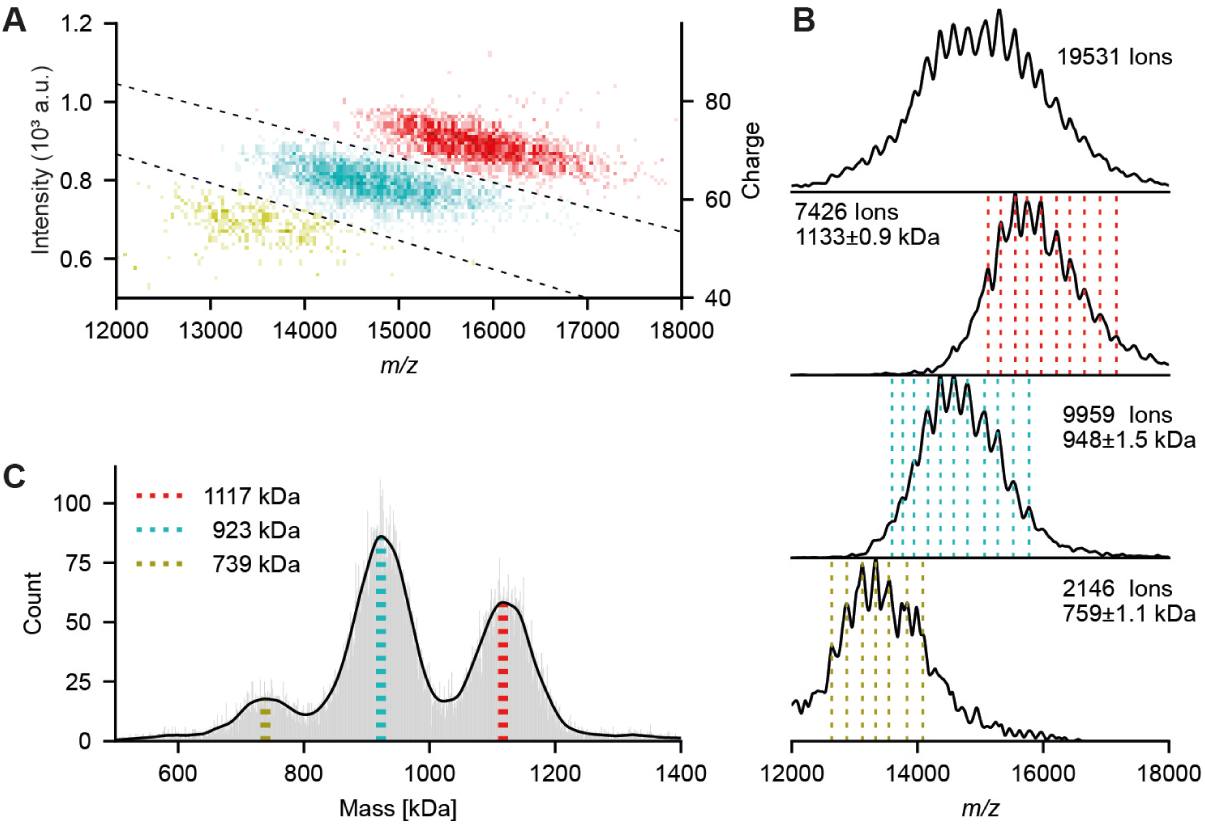
Resolving a complex mixture of IgM oligomers using single ion intensities. **A** Single particle CD-MS of IgM oligomers. Tetrameric, pentameric and hexameric species are coloured in red, cyan and green, respectively. Dotted lines indicate the margins used for the filtered subsets shown in **B**. **B** m/z histogram of single particle centroids showing the extensively overlapping charge state distributions for the three IgM oligomeric states (top). Below are shown the filtered subsets used for the charge state assignments. Dotted lines indicate top positions used for assignment and illustrate overlapping charge state series. **C** Mass histograms, calculated from ion intensities, revealing the distribution and masses of the three co-occurring species. The most abundant masses are indicated with vertical lines (same colour code as in **A** and **B**) and are in close agreement with the masses determined by using conventional charge assignments as depicted in **B**.

Using the intensity dimension, signals for each species could be clearly separated and intensity thresholds could be used as filtering criteria for each oligomeric species. The resulting *m/z* histograms shown in Fig. 5B can now be used for conventional charge assignment and quantification based on ion counts. The resulting determined molecular masses of 759 kDa, 948 kDa and 1,133 kDa correspond to the 4-mer, 5-mer and 6-mer and a monomer IgM mass of 189 kDa. Predicting charge and mass solely based on single particle intensity yields very similar masses for the 4-mer, 5-mer and 6-mer and a monomer mass of 185 kDa (see Fig. 5C). Both monomer masses are in accordance with the IgM monomer backbone mass of 173 kDa decorated with glycans on the six putative glycosylation sites.

### Single particle charge detection MS of megadalton virus assemblies

The paramount application for single particle CD-MS is in the analysis of large heterogeneous complexes of which charge states cannot be resolved in conventional native MS experiments. Similar to Pierson *et al*, AAV was used as a test case as its capsid is composed of 60 copies of three different building blocks (*i.e.* VP1, VP2 and VP3), assembled in varying stoichiometries, yielding a wide distribution of capsid masses ^30,41^. Genome-loaded AAV particles are even more heterogeneous, as the capsids package a mixture of sense and antisense ssDNA. To illustrate the challenges of native MS on AAV and the relative gains from single particle CD-MS, we measured AAV serotype 8 (AAV8) particles with a reported average 1:1:10 (VP1:VP2:VP3) ratio and loaded with a 3.8 kB genome encoding green fluorescent protein (GFP).

These AAV particles are so heterogeneous that no charge state assignment can be made in a conventional native MS spectrum (see Fig. S3). Following previous studies, mass can in this case only be roughly estimated from the average *m/z* position of the signal, assuming normal charging behavior for globular protein complexes based on empirical findings ^15,16^. With this approach we estimated the masses of the empty particle and its GFP genome filled counterpart at 3.8 and 5.8 MDa, respectively. Although the mass of the empty capsid is in accordance with the expected mass, this approach overestimates the mass of the genome-filled particle by about 1 MDa. This highlights the need for a more direct estimate of charge states by single particle CD-MS.

A single particle CD-MS analysis of AAV8 is presented in Fig. 6. We were able to resolve distinct distributions for the empty and packaged AAV8 particles. Strikingly, this analysis revealed that the empty and packaged AAV particles obtain a very similar number of charges during the native ESI, despite the filled particles having a 25% higher mass. The mass of the empty particle determined by CD-MS is 3,740 kDa, deviating by +11 kDa, or +0.3% of the expected mass. The mass of the genome loaded particles was calculated to be 4,910 kDa, yielding a genome mass of 1,170 kDa compared to the expected mass of 1,243 kDa, thereby confirming the packaging of the complete genome without substantial defects or degradation. The reported mass deviations are well within the error margin of ∼1% for MDa particles as discussed above.

**Figure 6:**
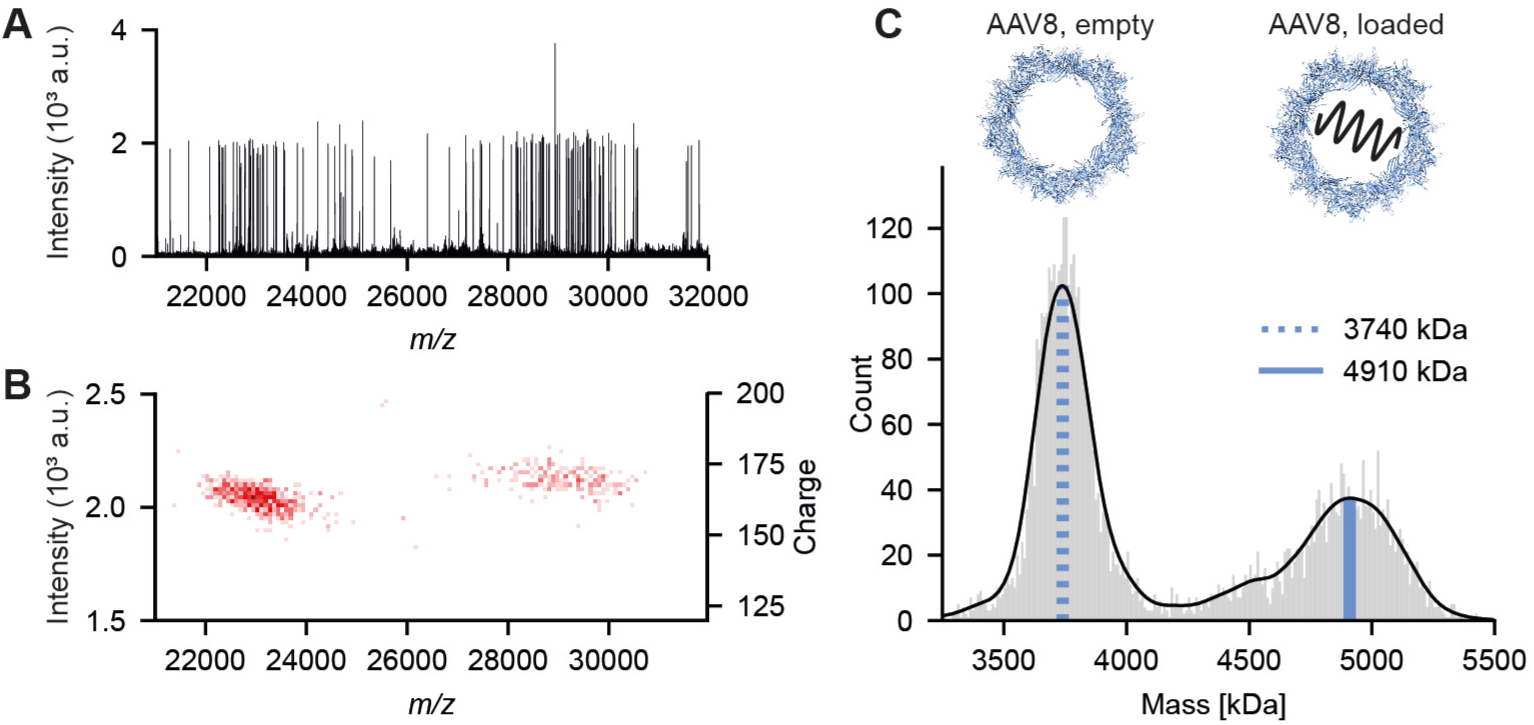
Single particle CD-MS of co-occurring empty and genome loaded AAV8 particles. **A** Individual scan of single particles for a mixture of empty and genome loaded AAV8 capsids. **B** 2D histogram of filtered single particles centroids collected over several minutes. **C** Mass histogram for AAV8 particles directly calculated from single ion intensities. Blue lines indicate top masses of the empty AAV8 (dotted) and the loaded AAV8 (solid) particle.

## Discussion

We demonstrate the capabilities of a commercial Orbitrap mass spectrometer to directly derive the charge state of particles based on single ion measurements. This overcomes a major bottleneck in native mass spectrometry, which normally requires charge-state resolved signals to assign charges and thus masses from *m/z* spectra. The current Orbitrap-based instruments are sufficiently sensitive to measure thousands of single particles in a matter of minutes, whereby the relative accuracy increases linearly with mass, making the technique even better suited for very large assemblies. Moreover, single particle CD-MS on Orbitrap MS is fully compatible with all its existing tandem MS and ion fragmentation capabilities. We foresee a thriving and broad range of applications for Orbitrap-based CD-MS, especially in analyzing complex oligomeric mixtures including amyloid fibrils, gene delivery vectors, highly glycosylated biotherapeutics and membrane protein complexes.

## Methods

### Native MS

Purified proteins were supplied from various sources: *E. coli* 70S Ribosome particles were purchased from Sigma-Aldrich, Trastuzimab and IgG1-RGY samples were provided by the team of Janine Schuurman at Genmab (Utrecht, NL), the IgM sample was provided by Suzan Rooijakkers (Medical Microbiology, UMCU, Utrecht, NL), the AaLS-neg nanocontainer sample was provided by the group of Don Hilvert (ETH Zurich) and the AAV8 particles were provided by Mavis Agbandje-McKenna (University of Florida).

The FHV and ribosomes samples were prepared and buffer exchanged as already described previously ^12^. All other purified protein samples were buffer exchanged to aqueous ammonium acetate (150 mM, pH 7.5) with several concentration/dilution rounds using Vivaspin Centrifugal concentrators (9,000xg, 4°C). An aliquot of 1–2 μl was loaded into gold-coated borosilicate capillaries 467 (prepared in-house) for nano-electrospray ionization. Samples were analysed on a standard commercial Q Exactive UHMR instrument (Thermo Fisher Scientific, Bremen, Germany)^12,42^. For conventional mass spectra ions optics were optimized for maximum ion transmission. For single particle acquisitions, ion transmission was attenuated by diluting the sample, reducing the collision gas pressures and reducing the injection time. Single particle datasets were acquired using either 512 or 1,024 ms set transients for 10 to 60 minutes.

### Single particle data pre-processing

Single particle data was first converted to mzXML files using the vendor peak picking algorithm (MSConvert)^43^. In order to get comparable intensity values throughout different experiments all ions intensities were multiplied by their injection time in seconds. Dephased ion signals were removed by applying an *m/z*-threshold for adjacent centroids above a certain absolute intensity value of 50. The applied *m/z* threshold depended on the *m/z* region the ions populated and we typically used five times the FWHM of the single particles peaks. See Supplementary Figure 4 for an overview of the workflow. A python class and an exemplary script for single particle data processing is available from the authors upon request.

### Charge vs. Intensity regression model

Pre-processed single ions datasets were filtered in the *m/z* and intensity dimension for the regions of interest. Centroid m/z-positions were then binned to obtain conventional mass spectra, which were used for conventional charge assignment. Single particle centroids were selected for each charge state based on their *m/z* position from the charge assignment. For each charge a kernel density estimation was performed in the intensity domain and peak intensity as well as FWHM of the distribution were extracted. From each set of intensities assigned to a certain charge, 200 samples were drawn randomly within two times the FWHM of the top intensity. All sampled ion intensities and their charges were subjected to linear regression. See Supplementary Figure 5 for an overview of the workflow.

### Intensity based charge and mass prediction

Pre-processed and filtered centroid intensities were converted into charge with the previously established regression model allowing non-integer values. Masses were then calculated from their *m/z* position using the equation: m = *m/z*\*z-z. Masses were subjected to a kernel density estimation and most abundant masses were extracted.

### Mass prediction of AAV based on *m/z position*

To estimate the mass from the *m/z* position, we fitted 76 empirical determined masses and their corresponding *m/z* positions to the equation Mass [kDa] = A * *m/z*^B^ as reported in previous publications ^15,16^. The resulting formula Mass [kDa] = 1.63 * 10^−6^ * *m/z*^2.14^ was used to estimate the mass based on the peak *m/z* position for the unresolved empty and genome loaded AAV8 particle from a spectrum recorded on a QToF instrument.

## Acknowledgments

We greatly appreciate that several collaborators provided samples to us that we used in the work presented here. We acknowledge the Don Hilvert group at the ETH Zurich (Switzerland) for providing the AaLS-neg samples, the Janine Schuurman group at Genmab (The Netherlands) for providing the mutant IgG samples, the Suzan Rooijakkers group at the Medical Microbiology Department, University Medical Center Utrecht (The Netherlands), in particular Piet Aerts and Carla Gosselaar-de Haas, for providing the IgM samples, and the Andrew Routh group at the University of Texas Medical Branch (USA), in particular Elizabeth Jaworski, for providing the FHV samples. This work was supported by the Netherlands Organization for Scientific Research (NWO) through the Spinoza Award, SPI.2017.028, to AJRH. Additional support came through the European Union Horizon 2020 INFRAIA project Epic-XS (Project 823839).

## Author Contributions

T.P.W., J.S. and A.J.R.H. conceived of the project, designed the experiments and wrote the paper. A.B. and M.A.-M. prepared the AAV samples. A.A.M. advised on data acquisition and processing. T.P.W. performed all experiments and processed the data. J.S. and A.J.R.H. supervised the project. T.P.W., J.S. and A.J.R.H. analyzed the results. All authors discussed the results and edited the paper.

## Supplementary Materials

Supplementary Table S1

Supplementary Figures S1-S5

