## Supplementary Materials for "Resolving heterogeneous high-mass macromolecular machineries by Orbitrap-based single particle charge detection mass spectrometry"

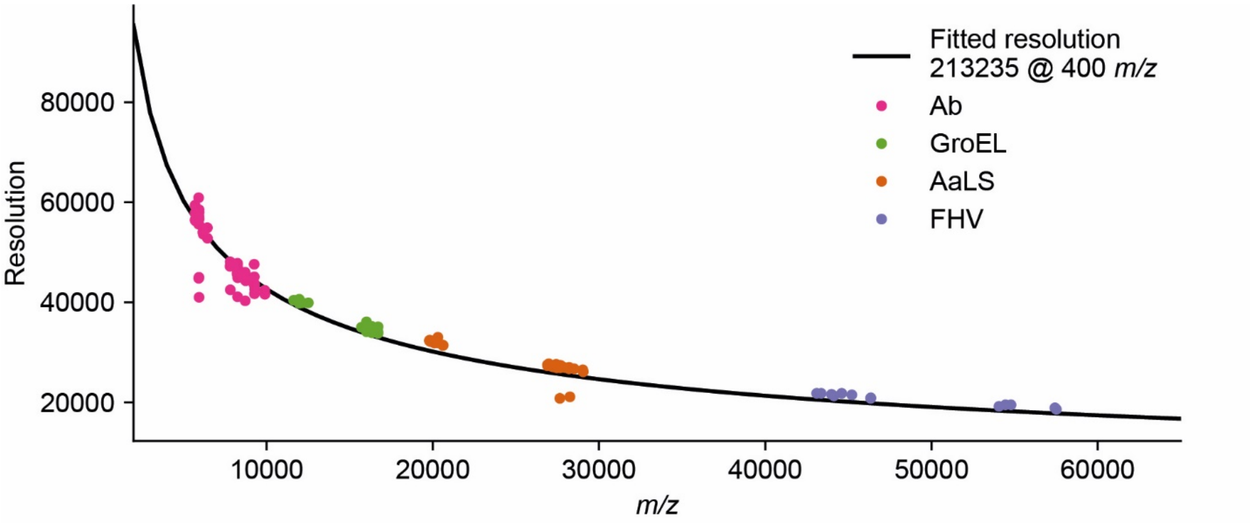


***Supplemental Figure 1: Single particle resolution*** - Extracted mass resolution for single particle events extracted from the data shown in Fig. 2 of the main text. Peak broadening causes some centroids to appear at slightly lower values than the theoretical resolution. Peak broadening can be caused by solvent loss during acquisition or if the ion does not survive over the set transient time. The theoretical resolution was obtained by fitting the dependency: *Resolution* ~ $m/z^{-1/2}$. The resulting resolution at 400 *m/z* is in close agreement with the reported values from the manufacturer for a 1,024 ms transient (21,3235 vs. 20,0000 at 400 *m/z*).

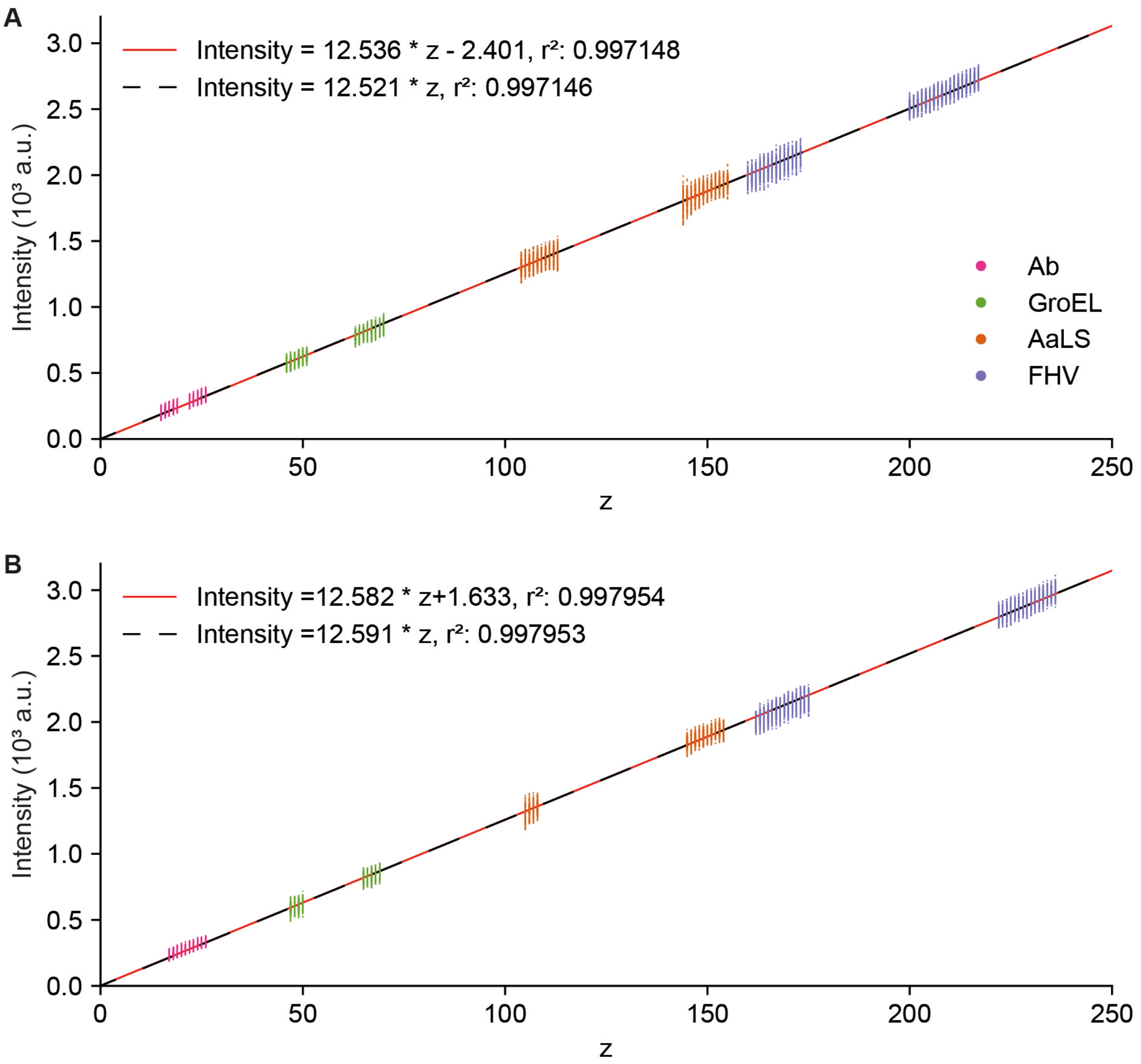


***Supplemental Figure 2: Linear regression models for Intensity vs. Charge correlation* - *A*** Linear regression models for the data shown in the main text. All data was recorded within three days. Data was fitted using a general linear model (red line) as well as one forced through the origin (black dashed line). Both regression models gave fits of similar quality ($\Delta r^2=2*10^{-6}$). The intersection at zero of the general regression model translates to less than 0.28 charges, hence we used for this work the one forced through the origin. **B** Technical replicate of the Intensity vs. Charge correlation shown in the main text and **A**. In contrast to **A**, data was acquired over a period of three months and at mixed resolution settings (512 ms and 1,024 ms). The maximum deviation between all shown regression models doesn't exceed 1.6 charges over the displayed range (1 to 250 charges).

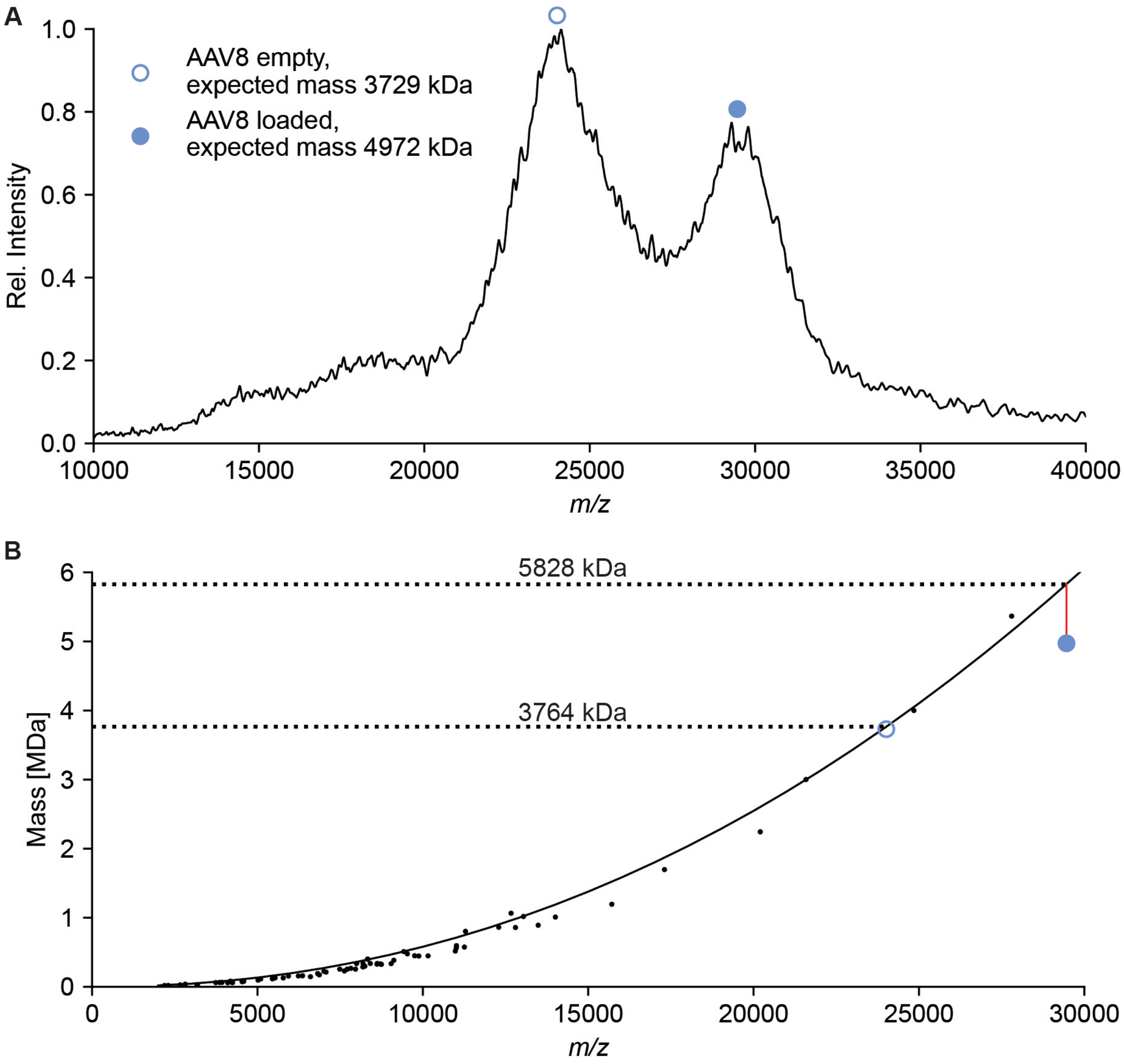


***Supplemental Figure 3: m/z position based mass prediction*** - **A** Mass spectrum of AAV8 recorded on an QToF mass spectrometer modified for high mass measurements. Peak positions for empty and genome loaded AAV8 are indicated with empty and solid blue circles (24,012 *m/z* and 29,459 *m/z*, respectively). **B** Empirical dependency between *m/z* position and mass for 76 globular proteins, measured and reported previously, fitted with an exponential function. The resulting formula $Mass[kDa] = 1.63 * 10^{-6} * m/z^{2.14}$ was used to calculate the mass for the empty and genome loaded AAV8 particle (3,764 kDa and 5,828 kDa respectively). The calculated mass for the empty AAV8 is in close agreement with the expected mass but the mass of the genome loaded particle deviates by ~1 MDa.

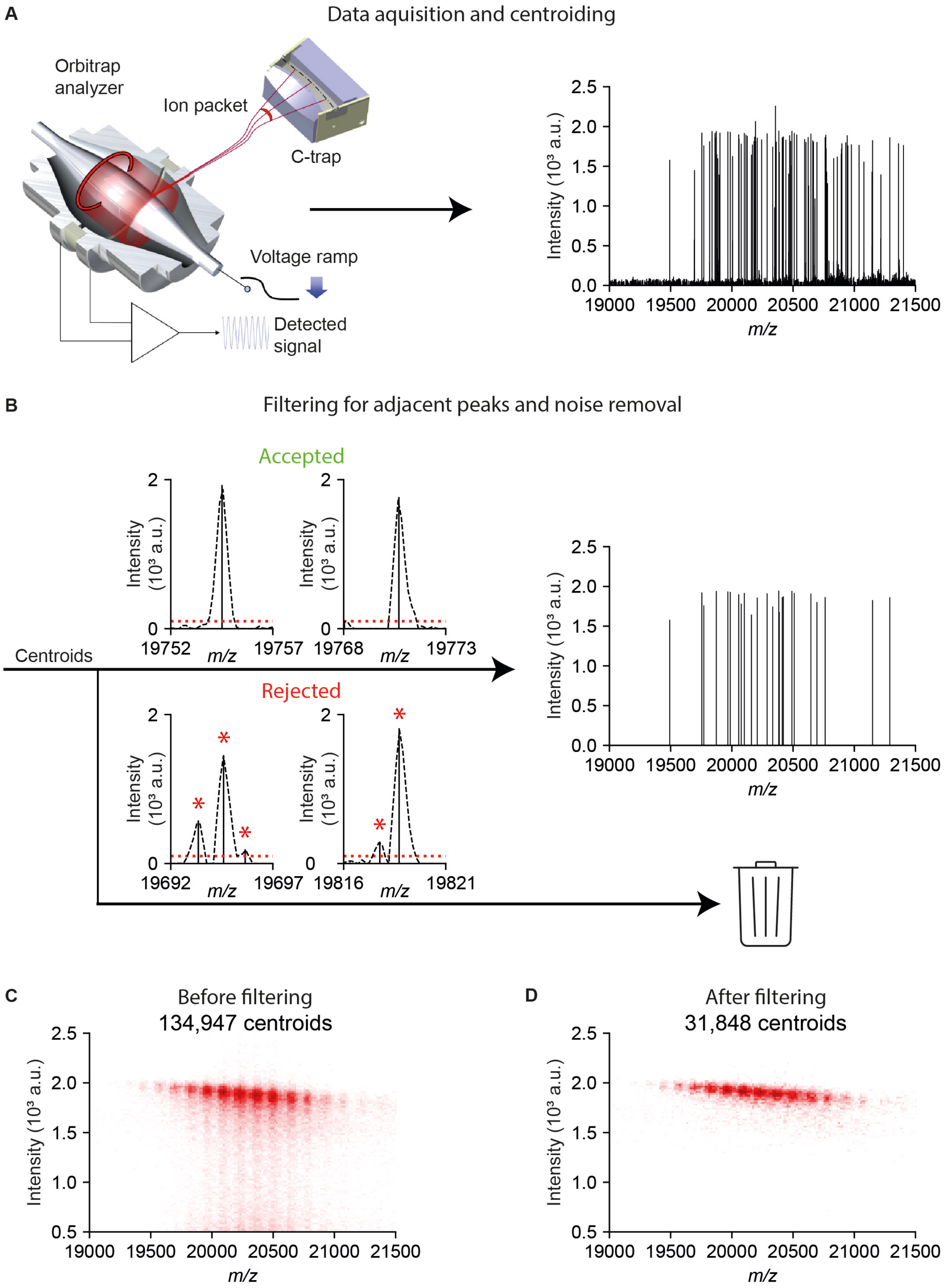


***Supplemental Figure 4: General workflow for single particle pre-processing*** - **A** Data was acquired on a QE-UHMR. For the data shown in this work, scans were collected for 10 to 30 minutes. After acquisition, .RAW files were centroided and converted into the mzXML format. **B** Centroids were subjected to filtering. We rejected all centroids which were below a certain intensity value as well as centroids which have an adjacent peak within a certain *m/z* threshold above the noise limit (See red asterisk in rejected traces). **C&D** Effect of filtering on AaLS-neg dataset. **C** 2D histogram of single particle centroids before filtering. Ion intensities trail toward lower values, which is mainly attributed to the presence of split peaks presumably resulting from dephasing and/or mass loss of particles during the transient. Both effects are expected to originate from collisions with background gas and/or metastable decay during detection. **D** Dataset after noise filtering and removal of adjacent peaks. Filtered single particle centroids are more confined in the intensity domain and don't show the trailing towards lower intensities anymore.

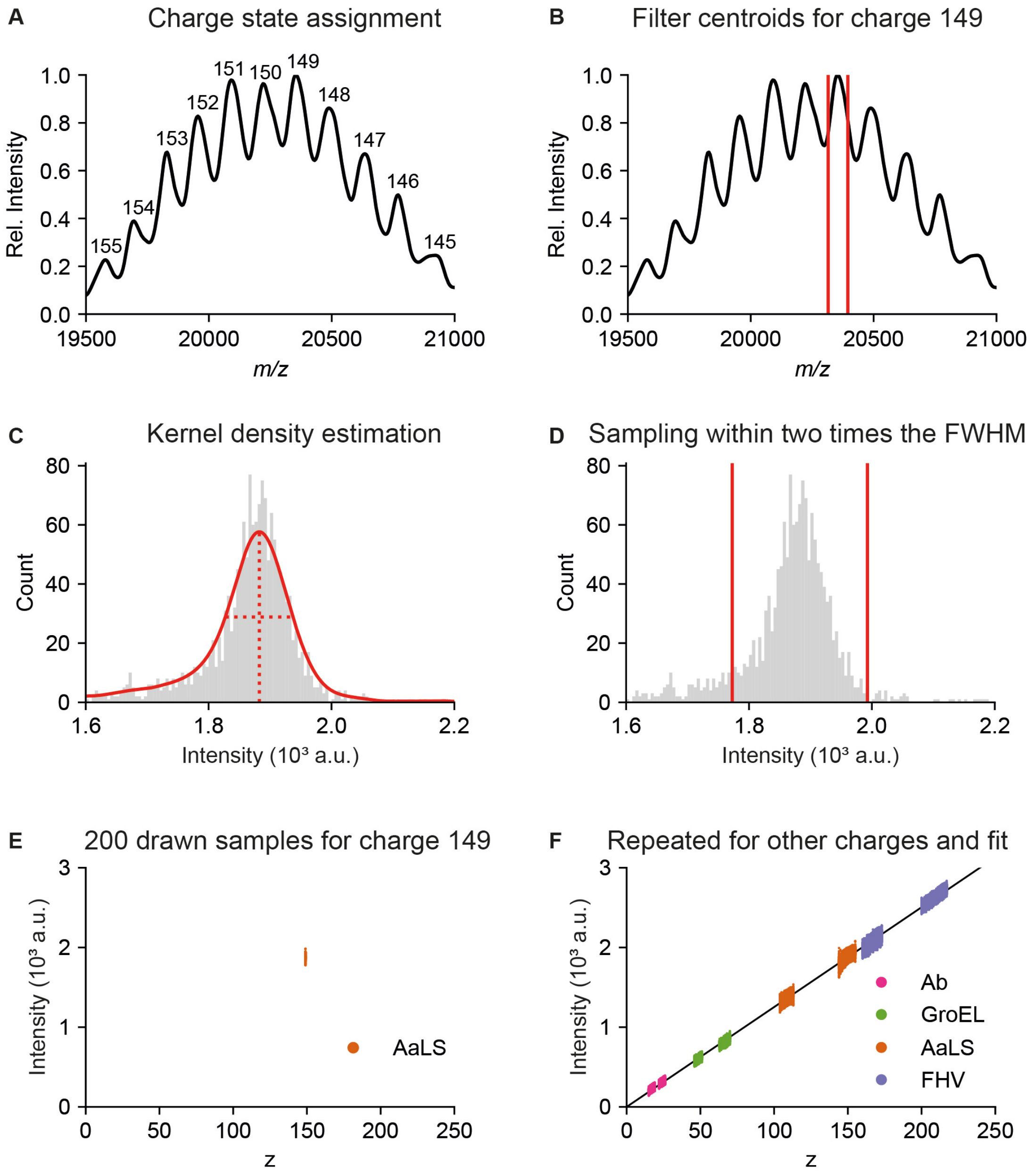


***Supplemental Figure 5: General procedure for establishing Intensity vs. Charge regression model shown for AaLS-neg, (z=149)*** - **A** Single particle centroids, preprocessed and filtered as described in Supplementary Figure 4, were binned in *m/z* and smoothed. The resulting mass spectrum was used for conventional charge sate assignment. **B** Centroids were filtered for charge state 149 by only allowing centroids within 80 Th of the top peak position to pass the filtering. **C** Filtered centroids were subjected to a kernel density estimation and the most abundant intensity as well as the FWHM of the distribution were extracted. **D** To obtain an equal weighted fit over all tested charges 200 random samples were drawn for each charge within two times the FWHM of the most abundant intensity. Assuming a normal distribution this contains more than 98

% of the centroids (FWHM = 2.35σ). **F** The same procedure was repeated for the other examined proteins and charges and used for establishing the linear regression model.

***Supplemental Table 1: Overview of expected and measured masses of proteins assemblies analysed in this study*** – Table shows on overview of the expected masses for the analysed protein assemblies as reported on previous studies. Masses were either calculated via the conventional charge states assignment strategy (CS) or by predicting the charge from its intensity via Orbitrap-based charge detection (CD). Mass deviations are expressed as percentage compared to the expected mass (ΔE) or compared to the mass calculated via the conventional charge states assignment strategy (ΔCS).

| Protein | TEAA [mM] | Expected [kDa] | Measured (CS) [kDa] | ΔE [%] | Measured (CD) [kDa] | ΔE [%] | ΔCS [%] |
|---|---|---|---|---|---|---|---|
| **Antibody** | - | 148[a] | 148 | 0.3 | 154 | 4.1 | 3.7 |
| | 25 | 148 [a] | 148 | 0.3 | 160 | 8.1 | 7.8 |
| **GroEL** | - | 801[1] | 801 | 0.0 | 786 | 1.9 | 1.9 |
| | 25 | 801[1] | 801 | 0.0 | 794 | 0.9 | 0.9 |
| **AaLS** | - | 3015[2] | 3032 | 0.6 | 3060 | 1.5 | 0.9 |
| | 25 | 3015[2] | 3047 | 1.1 | 3070 | 1.8 | 0.8 |
| **FHV** | - | 9307[3] | 9361 | 0.6 | 9400 | 1.0 | 0.4 |
| | 25 | 9307[3] | 9364 | 0.6 | 9260 | 0.5 | 1.1 |
| **IgM tetramer** | - | 745[b] | 759 | 1.9 | 739 | 0.8 | 2.6 |
| **IgM pentamer** | - | 931 [b] | 948 | 1.8 | 923 | 0.9 | 2.6 |
| **IgM hexamer** | - | 1117 [b] | 1133 | 1.4 | 1117 | 0.0 | 1.4 |
| **AAV8 empty** | - | 3729[4] | - | | 3740 | 0.3 | |
| **AAV8 loaded** | - | 4972 [c] | - | | 4910 | 1.2 | |

**a**: Average mass of the three main glycoforms (G0F/G0F, G0F/G1F and G0F/G2F) as reported in “Full characterization of heterogeneous antibody samples under denaturing and native conditions on the Q Exactive BioPharma mass spectrometer” by Thermo Fisher Scientific.

**b**: Masses of oligomeric species were calculated from the IgM backbone mass (173 kDa) plus 2.2 kDa per glycosylation site (six in total).

**c**: Genome mass was calculated for a 3.8 kB genome, approximating 327 Da/base.

1. Sobott, F. & Robinson, C. V. Characterising electrosprayed biomolecules using tandem-MS - The noncovalent GroEL chaperonin assembly. *Int. J. Mass Spectrom.* **236**, 25–32 (2004).

2. Sasaki, E. *et al.* Structure and assembly of scalable porous protein cages. *Nat. Commun.* **8**, 14663 (2017).

3. van de Waterbeemd, M. *et al.* High-fidelity mass analysis unveils heterogeneity in intact ribosomal particles. *Nat. Methods* 1–7 (2017). doi:10.1038/nmeth.4147

4. Pierson, E. E., Keifer, D. Z., Asokan, A. & Jarrold, M. F. Resolving Adeno-Associated Viral Particle Diversity with Charge Detection Mass Spectrometry. *Anal. Chem.* **88**, 6718–6725 (2016).
